# Mask, the *Drosophila* Ankyrin Repeat and KH domain-containing protein, regulates microtubule dynamics

**DOI:** 10.1101/2020.04.22.056051

**Authors:** Daniel Martinez, Mingwei Zhu, Jessie J. Guidry, Niles Majeste, Hui Mao, Sarah Yanofsky, Xiaolin Tian, Chunlai Wu

## Abstract

Proper regulation of microtubule (MT) dynamics is vital for essential cellular processes and neuronal activities, including axonal transport as well as synaptic growth and remodeling. Here we demonstrate that Mask negatively regulates MT stability and maintains a balanced dynamics of MT length and architecture in both fly larval muscles and motor neurons. In larval muscles, loss of *mask* increases MT length, and genetically altering *mask* levels modifies the Tau-induced MT fragmentation. In motor neurons, loss of *mask* function reduces the number of End-Binding Protein 1 (EB1)-positive MT plus-end structures in the axons and also results in overexpansion of the presynaptic terminal at larval neuromuscular junctions (NMJs). *mask* shows strong genetic interaction with *stathmin (stai)*, a neuronal modulator of MT dynamics, in regulation of axon transportation and synaptic terminal stability. Our structure/function analysis on Mask revealed that truncated Mask transgenes carrying only its N-terminal portion that contains the two Ankyrin repeats domains is able to rescue the MT-related *mask* loss-of-function defects in larval muscles and NMJs, suggesting an essential role of the Ankyrin repeats domains in mediating Mask’s MT stability-regulating function. Furthermore, we discovered that Mask negatively regulates the abundance of the microtubule-associated protein Jupiter in motor neuron axons, and that neuronal knocking down of *Jupiter* partially suppresses *mask* loss of function phenotypes at the larval NMJs. Together, our study identified Mask as a novel regulator for microtubule stability and dynamics.

**Author Summary:** Microtubules (MTs) are an essential part of the cellular cytoskeleton, providing the structural basis for critical cellular processes and functionality. A series of factors are required to orchestrate the assembly and disassembly of MTs. Here, we identified Mask as a novel regulator of MT dynamics in *Drosophila*. Mask shows prominent interplay with two important modulators of MT, Tau and Stathmin (Stai). These findings not only support the role of Mask as a novel microtubule regulator but also provide the foundation to explore future therapeutic strategies in mitigating deficits related to dysfunction of Tau and/or Stai, both of which are linked to human neurodegenerative disorders. Further analyses on Mask demonstrated that Jupiter’s localization to the MTs in the axons is negatively affected by Mask, and that reducing Jupiter level is able to partially suppress synaptic defects caused by *mask* mutant. Together, these data imply that Mask’s function in regulating MT dynamics requires Jupiter.

## Introduction

Terminally differentiated postmitotic cells such as neurons and muscles use their microtubule (MT) network not for cell division but rather as architectural components essential for their shape and unique cellular functions. In addition to supporting the structural integrity of the axons and the dynamic morphological changes of the dendritic in neurons, MTs also act as directional railways for transporting materials and organelles between the cell body and the synapses (1). MTs can undergo cycles of dynamic assembly and disassembly (labile state) or stay relatively stable in a cell type- and developmental stage-dependent manner (2). For example, MTs in postmitotic cells, such as neurons, are generally more stable than MTs in dividing cells. However, within a developing neuron, MTs at the axon growth cone are much more labile than MTs near the soma (3), and even in individual axons, MT network consists of domains that differ in their stability (4). Recent studies suggest that both the stable and labile pools of microtubules play essential roles for normal neuronal function. Furthermore, the spacing among MTs and distance between MT ends have both been shown to be critical for normal axonal transport (5, 6). Therefore, striking a balance in MT dynamic is of vital importance to maintaining MT-mediated cellular functions.

Many proteins and pathways have been identified as potential regulators of MT stability (2, 7). Among the major proteins controlling MT stability, Stathmin/SCG10 (superior cervical ganglion-10 protein) and Tau/MAPT are both MT-binding proteins which regulate multiple aspects of MT stability, including growth and shrinkage, as well as the transition between catastrophe and rescue. Both Stathmin and Tau are associated with diverse models of neurodegeneration, axon transport defects, and cancer (8–11). While *in vitro* studies of Stathmin-related proteins in mammals suggest that Stathmin promotes destabilization of MTs, studies of *stathmin (stai)* in fly neuromuscular junction (NMJs) showed that it is required for MT stabilization, as well as axon transport and NMJ stability (12, 13). The results of the *in vivo* studies in mammals is consistent with the fly data in that *stathmin* knockout mice exhibit age-dependent axonopathy in both the central and peripheral nervous systems, and they exhibit defective motor axon outgrowth and regeneration (14, 15). Tau, on the other hand, plays a multifaceted role in cell survival signaling. Loss of Tau function or high levels of hyperphosphorylated Tau disrupts MT stability, leading to axonal transportation defects in motor neurons and microtubule breakdown in larval muscles (16). Additionally, hyperphosphorylated Tau aggregates form inclusion bodies that are associated with a variety of disorders collectively referred to as tauopathies, including Alzheimer’s disease(17). In animal models, such as rodents and fruit flies, overexpression of human Tau in the neuronal tissues leads to progressive neurodegeneration (18).

Mask is a 4001-amino-acid protein with several functional domains. It consists of two Ankyrin Repeats, one Nuclear Export Signal (NES), one Nuclear Localization Signal (NLS), and a C-terminal KH domain. The two Ankyrin repeat domains containing 15 and 10 tandem Ankyrin repeats likely coordinately facilitate the ability of Mask to associate with other proteins, according to the well-documented involvement of the Ankyrin domains in mediating protein-protein interactions in eukaryotic cells (19). The NES and NLS motifs may be required for shuttling Mask protein in and out of the nucleus, which is essential for its interaction with the Hippo pathway effector Yorkie/YAP in mitotic cells (20, 21). The KH domain is an evolutionarily conserved motif that is about 70 amino acids long, and it was first identified in the human heterogeneous nuclear ribonucleoprotein K (22). KH domains bind RNA or single-stranded DNA (ssDNA) and are found in proteins involved in transcription, translation and mRNA stability regulation (23, 24). Mask has been linked to several signaling pathways and different cellular processes in mitotic cells-it regulates cell proliferation during development (25); it is a component of the centrosome and nuclear matrix (26, 27) and a co-transcription factor of the Hippo pathway (20, 21). The human homolog of Mask, ANKHD1, is expressed at relatively high levels in acute leukemia cells (28), multiple myeloma cells (29) and prostate cancer cells (30). In the cancer cells, ANKHD1 is shown to be able to suppress p21 (31) and Stathmin activity (32). Regardless the attentions that has been casted on its function in mitotic cells, the functions of Mask and its vertebrate homolog in post-mitotic cells, including neurons and muscle cells, is largely unknown. Our previous studies on Mask demonstrated that Mask regulates mitochondrial morphology (33) and promotes autophagy (34) in larval muscles. Here, we show that Mask is required for balanced MT stability in muscle and neurons, two post-mitotic cell types, and that Mask genetically interacts with two MT-interacting proteins, Tau and Stathmin, whose dysfunctions are linked with human diseases. Furthermore, the abundance of Jupiter, a microtubule-associated protein, is reversely related to Mask levels in the motor neuron axons. Knocking down *Jupiter* in the neurons partially suppresses the morphological defects caused by *mask* loss-of-function at the larval NMJs. Together, our studies demonstrated a novel function of Mask in regulating microtubule dynamics.

## Results

### Mask negatively regulates MT length in larval muscles

Our previous studies of the putative scaffolding protein Mask demonstrated that overexpressing Mask ameliorate the degeneration of photoreceptors caused by overexpressing Tau in adult fly eyes (34). This finding prompted us to explore further potential interplays between Mask and Tau in the context of microtubule (MT) morphology, given the fact that Tau is a well-studied MT-binding protein. We first examined MT morphology in *mask* loss-of-function mutants. Interestingly, we found that, in the larval body wall muscles, the MTs in the *mask* null mutants are substantially longer than those in the wild type control, particularly in the area surrounding the muscle nuclei (Fig. 1AB). Such a phenotype is fully rescued by introducing the UAS-Mask transgene back to the *mask* mutant larval muscles (Fig. 1AB). To further confirm this finding, we introduced *UAS-mask* RNAi to the wild type larval muscle and found that *mask* knockdown also moderately increased muscular MT length (Fig. 1AB). These data suggested that one of the normal functions of Mask in larval muscles is to restrain MT length.

**Figure 1.**
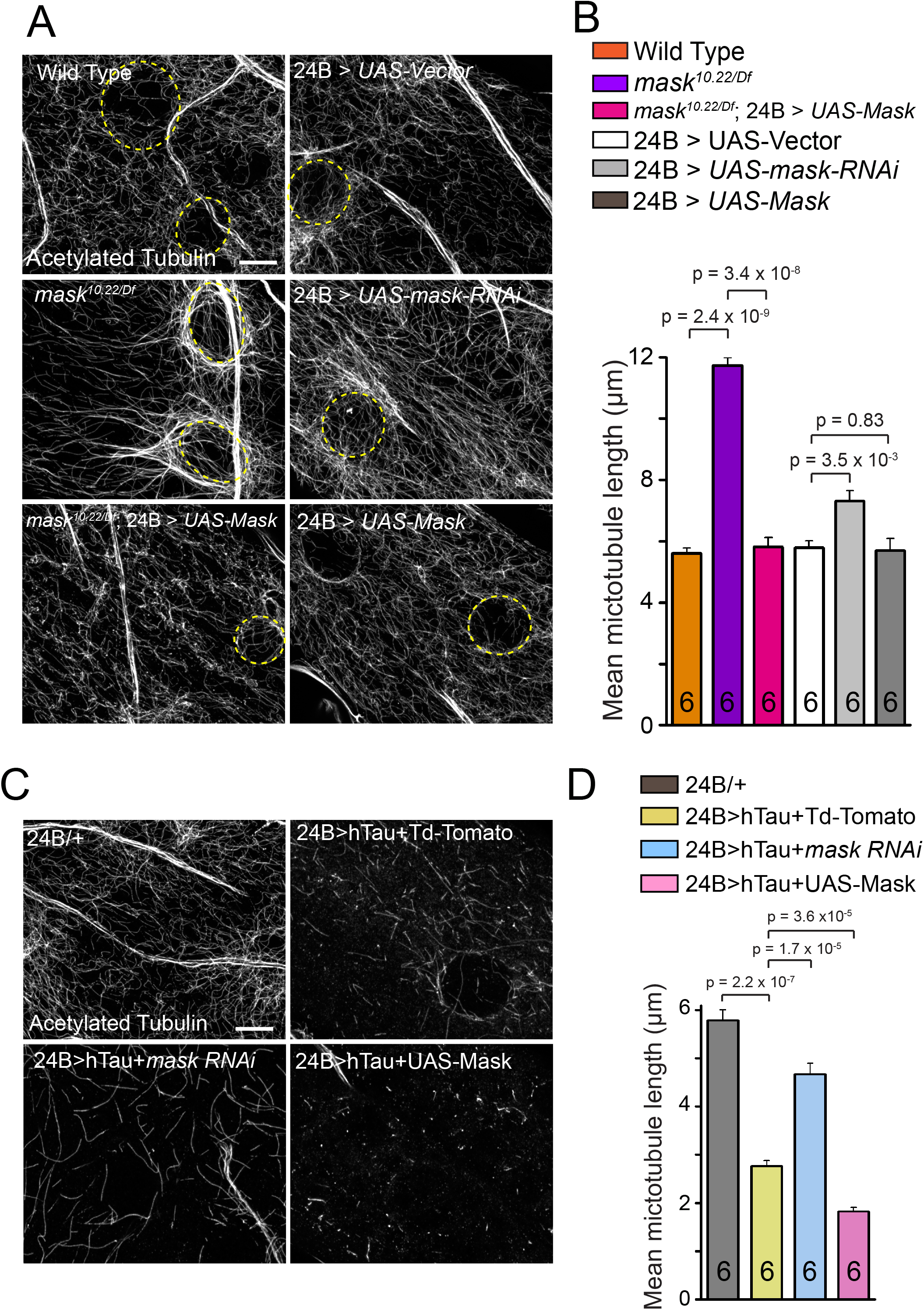
*mask* negatively regulates microtubule stability in larval muscle and enhances Tau-induced MT fragmentation. (**A**) Representative confocal images of microtubule (MT) in muscle 6 of wild type, *mask* null *(mask^10.22/Df^)*, rescue of *mask* null with a 24B (muscle-specific)-Gal4-driven UAS-Mask transgene, 24B -Gal4-driven UAS-Vector, *UAS-mask* RNAi, or UAS-Mask.. MTs are immunostained with an anti-Acetylated Tubulin antibody. Yellow dash lines denote the edge of muscle nuclei. Scale bar: 5 μm. (**B**) Quantification of average MT lengths. (**C**) Representative confocal images of MT in muscle 6 of 24B-Gal4/+, 24B-Gal4-driven UAS-human Tau (hTau) with UAS-Td-Tomato, *UAS-mask* RNAi, or UAS-Mask. MTs are immunostained with an anti-Acetylated Tubulin antibody. Scale bar: 5 μm. (**D**) Quantification of average MT lengths.

### Loss of *mask* function suppresses, while upregulation of Mask enhances Tau-induced MT breakdown and toxicity in fly muscles

Next, we examined the interplay between Mask and Tau in regulating the MT network in larval muscles. Overexpression of human Tau together with UAS-tdTomato (control) in fly muscles causes severe destruction of the MT network, and the affected MTs becomes short and punctate (Fig. 1 C), which is consistent with a previously reported study on human Tau overexpression in fly muscles (16). *mask* RNAi in the muscle, with its effects to increase MT length, was able to substantially reverse the shortened MT length caused by Tau-overexpression. Overexpressing Mask alone in wild type muscles, on the other hand, although alone does not lead to obvious reduction in the MT length in larval body wall muscles (Fig. 1AB), when co-expressed with Tau further devastates the MT network – MT almost only exists in bright rod-shaped puncta (Fig. 1C,D), suggesting the ability of Mask-overexpression to impact MT morphology under the sensitized background. To test whether Mask knockdown or overexpression alter the expression levels of human Tau, which in turn modifies Tau-induced MT defects, we examined the expression levels of human Tau under the conditions of up- or down-regulation of Mask. Expression of human Tau in the muscles is increased when it is co-expressed with UAS-Mask RNAi, suggested that the suppression effects are not induced by reducing Tau expression level by Mask knockdown (Fig. S1). Similarly, levels of human Tau expression in the muscle are decreased when it is co-expressed with UAS-Mask, therefore, the enhancement of the MT fragmentation is not a result of further elevation of Tau in the muscles (Fig. S1). These data are consistent with our previous finding that Mask actively promotes autophagy and degradation of aggregation-prone proteins in larval muscles (34).

Our morphological and genetic analyses suggested that the normal function of *mask* is to restrain MT length. To further confirm this notion, we used additional biochemical analysis to examine the overall distribution of MT length in the cell lysates of wild type or *mask* knockdown larval muscles. We found that reducing Mask levels does not alter the total Tubulin concentration in muscle homogenates (Fig. 2A). In the presence of 100 μM taxol (a potent concentration that promotes microtubule polymerization), Tubulin proteins in both control and mask *knockdown* lysates polymerize and fractionate exclusively in the pellet after ultracentrifugation (Fig. 2A). In the presence of 0.1 μM taxol (a concentration that reverses a steady state of MT dynamic status), *mask* RNAi lysate contains more Tubulin proteins that fractionate with pellet fraction (Fig. 2 A, B) compared to the control condition, suggesting that loss of Mask activity in muscles results in a MT network that is comprised of longer MTs.

**Figure 2.**
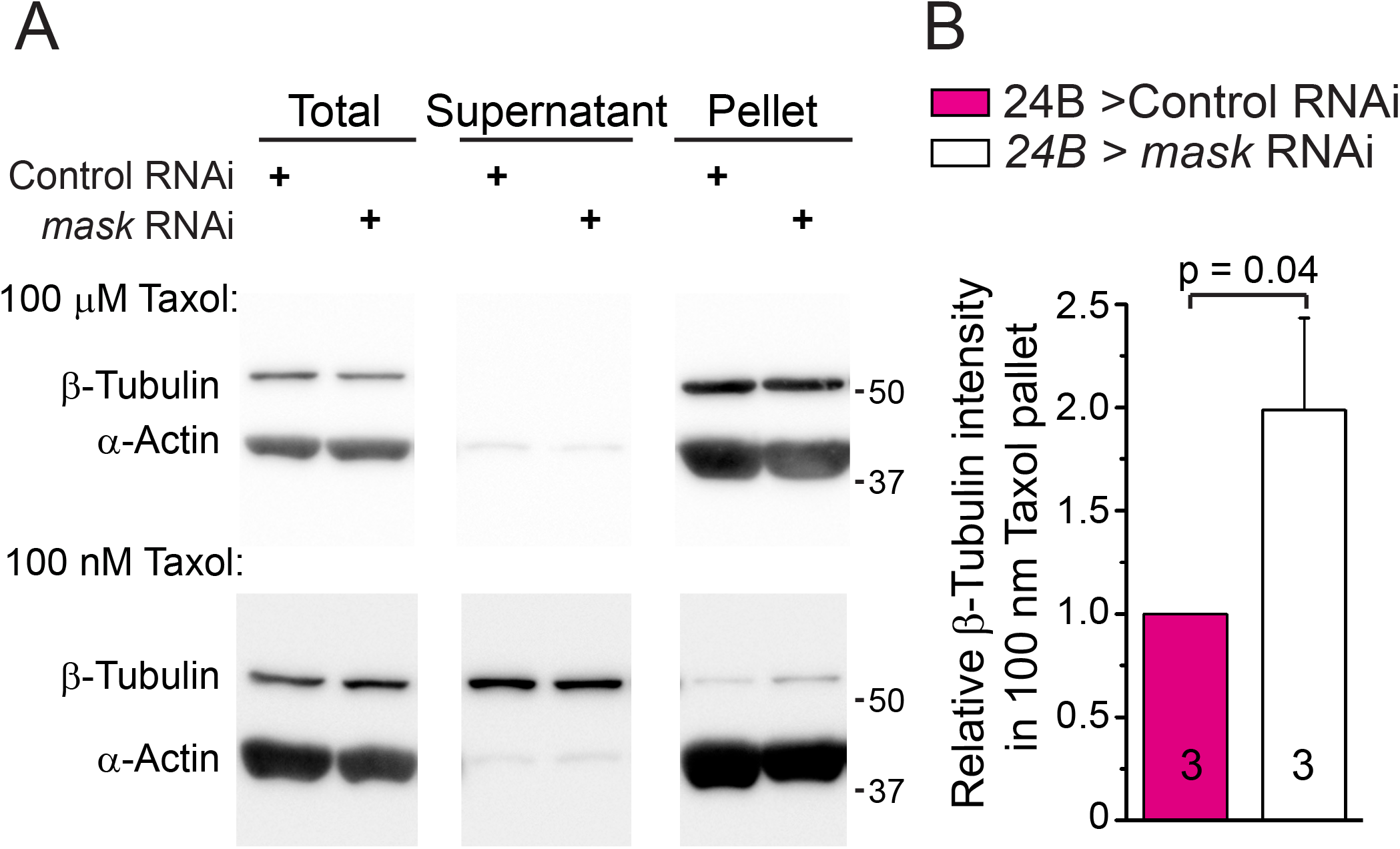
*mask* knockdown increase formation of longer MTs in fly larval muscles. **(A)** Western analysis of total proteins and ultracentrifugal fractions (supernatant and pellet) in larval muscle lysates from wild type control or *mask* knockdown (24B-Gal4-driven *mask* RNAi). Lysates were treated with either 100 μm or 100 nm Taxol. Anti-beta-Tubulin and anti-alpha-Actin blots were performed on total and ultracentrifugal fractions of both lysates. **(B)** Quantification of the relative levels of Tubulin fractionated in the pellet fraction in the lysate treated by 100 nm Taxol. The levels of Tubulin in pellet were normalized to the levels of total Tubulin. n = 3 trials.

### Mask regulates presynaptic terminal growth in neuromuscular junctions

The prominent protective effects of Mask in the photoreceptors (35) also prompted us to analyzed the neuronal functions of *mask* using the fly larval NMJ as a model synapse. We found that *mask* null mutant (*mask^10.22/Df^*) NMJs show expanded presynaptic terminal growth reflected by an increased number of bouton, synaptic span, and number of branching points (Fig. 3AB). Such a morphological defect is due to loss-of-function of *mask* in the presynaptic motor neurons as pan-neuronal or ubiquitous expression, but not muscle (postsynaptic) expression, of UAS-Mask rescues the NMJ overgrowth phenotype in *mask* null mutants (Fig. 3AB). Furthermore, neuronal knockdown of *mask* using a *UAS-mask* RNAi causes similar NMJ expansion as *mask* mutants (Fig. 3AB). These data demonstrate that Mask regulates normal NMJ expansion in a cell-autonomous manner in neurons. Given that microtubule stability has a direct impact on synaptic size and morphology(36), we then investigated whether the neuronal regulation on the presynaptic expansion by Mask is connected with its function in regulating MT dynamics. We first analyzed MT dynamics in the segmental nerve using the MT plus-end tracking protein End-Binding Protein 1 (EB1) (37) as the marker to assess the abundance of the MT plus-end structures. In motor neuron axons, EB1-GFP proteins exhibit two types of localization patterns - a uniform distribution along the axons, and bright comets scattered within the axons- the signals that represent the accumulation of the end-binding protein at the MT plus-ends. We found that the numbers of the EB1-GFP comets are significantly reduced by neuronal knockdown of *mask* when compared to the controls (Fig. 3CD). These results together suggest that loss of *mask* leads to formation of fewer plus-end structures, and that MTs present in the axons are likely structurally longer and less dynamic.

**Figure 3.**
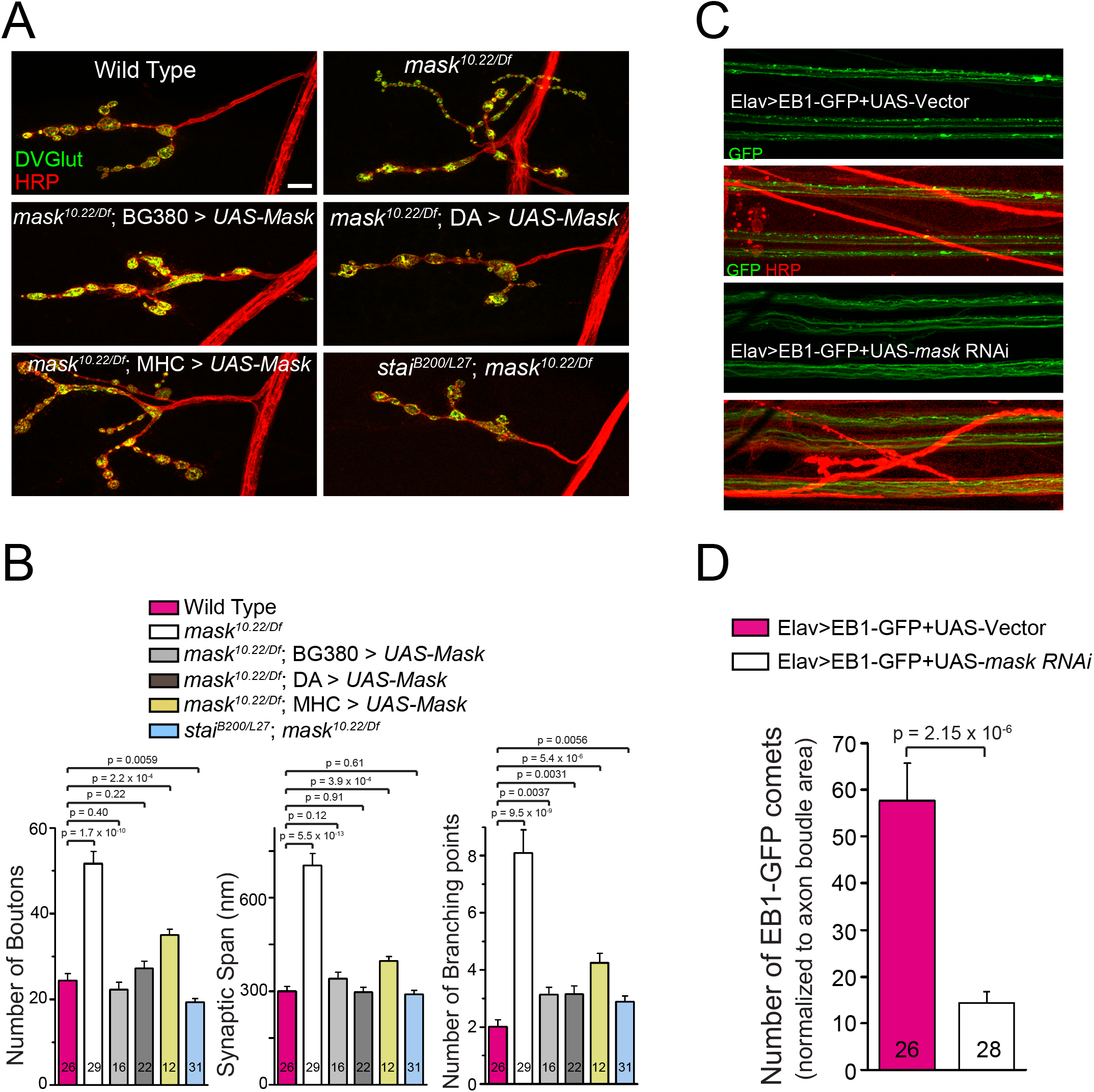
Mask promotes normal NMJ terminal growth by regulating motor neuron microtubule stability. (**A**) Representative confocal images of larval muscle 4 NMJs in wild type, *mask* null (*mask^10.22/Df^), mask* null rescued with a UAS-Mask transgene driven by pan-neuron (BG380), ubiquitous (DA) or muscle (MHC) Gal4 drivers, and *stathmin/mask* double mutant (*stai^B200/L27^; mask^10.22/Df^*). Scale bar: 10 μm. (**B**) Quantification of the number of boutons, synaptic span and the number of branching points at the Muscle 4 NMJs. Each data point was normalized to the size of the imaged muscle 4. (**C**) Representative confocal images of native GFP fluorescence plus a very brief staining of Cy3-HRP (see method section) of segmental nerves in 3^rd^ instar larvae containing Elav-driven UAS-EB1-GFP with UAS-vector or with *UAS-mask* RNAi. Scale bar: 20 μm. (**D**) Quantification of average number of EB1-GFP comets normalized to nerve area.

### *mask* and *stathmin* genetically interact with each other to regulate morphology and structural stability of NMJs

To further investigate Mask’s action in regulating MT elongation in neurons, we tested genetic interaction between *mask* and *stathmin. stathmin* is a well-established modulator of MT stability in neurons. Loss of *stathmin* causes severe destabilization of MT in motor neurons, resulting in axonal transport defects, especially in the posterior segments, as well as premature loss of presynaptic structure at the nerve terminals (named the “footprint” phenotype in *Drosophila* neuromuscular junctions) (12, 13). Mammalian studies showed that ANKHD1, the human homolog of *mask*, regulates the activity of Stathmin 1 in leukemia cells(32), suggesting a possible link between the two genes in neurons. We found that the presynaptic NMJ expansion observed in *mask* mutant NMJs is completely suppressed in the *stai/mask* double mutant NMJs (Fig. 3AB). Similarly, the NMJ expansion phenotype induced by neuronal knockdown (RNAi) of *mask* can also be partially suppressed by a heterozygous *stai* mutant, and completely suppressed by the homozygous *stai* mutant (Fig. 4AB), suggesting a strong genetic interaction between the two genes. In addition, *mask* loss of function can also reversely suppress neuronal defects caused by loss of function of *stai*. Both the footprint phenotype and the axonal transport defect of *stai* mutants are partially suppressed by the neuronal knockdown of *mask*, and completely suppressed by the *mask* null mutant (Fig. 4CD), suggesting that the enhanced MT stability by *mask* loss of function is able to compensate the impaired MT stability caused by loss of *stai*. These data provides strong evidences to support the notion that *mask* and *stai* exert opposite effects on MTs and these effects antagonize each other in regard to MT stability. The strong genetic interactions between *mask* and *stai* further suggest that *mask* regulates synaptic morphology and formation at the larval NMJs via its control over MT stability.

**Figure 4.**
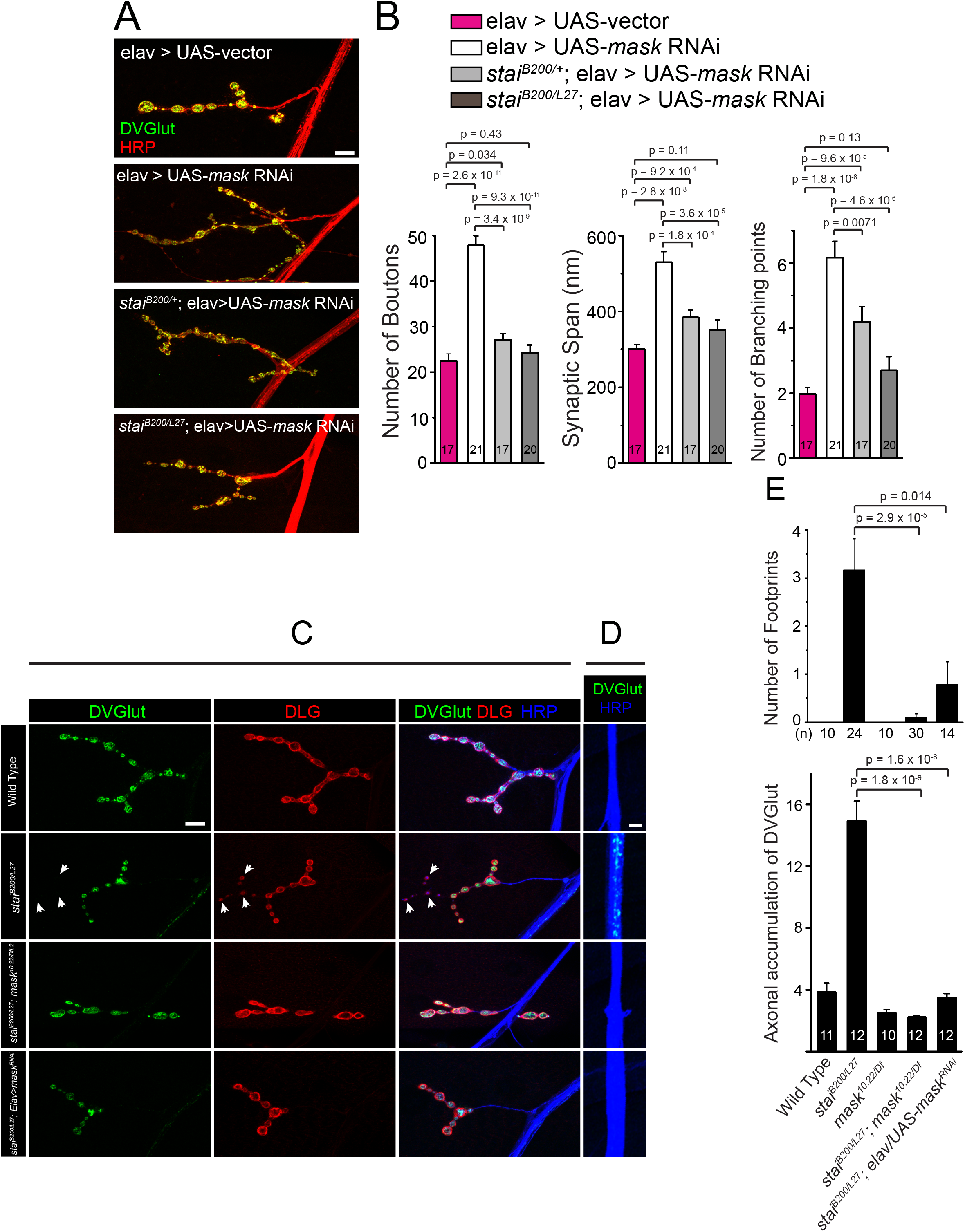
*mask* and *stai* reciprocally suppress each other. (**A, B**) Loss of *stai* function suppresses the synaptic terminal over-expansion caused by *mask* loss-of-function in a dose-dependent manner. (**A**) Representative confocal images of muscle 4 NMJs in larvae with Elav-driven UAS-vector, Elav-driven *UAS-mask* RNAi in wild type background, *stai* heterozygous (*stai^B200/+^*) or *stai* homozygous (*stai^B200/L27^*)mutant backgrounds. Scale bar, 10 μm. (**B**) quantification of the number of boutons, synaptic span and the number of branching points at the muscle 4 NMJs. Each data point was normalized to the size of the imaged muscle 4. NMJs were immunostained with anti-HRP (red) and anti-DVGlut (green) antibodies. (**C, D, E**) Loss of *mask* function in neurons suppresses *stai* mutant defects in NMJ development and axonal transport. (**C, D**) Representative confocal images of muscle 4 NMJs (C) and lateral nerve bundles (D) in wild type, *stai (stai^B200/L27^), stai/mask* double mutant (*stai^B200/L27^;mask^10.22/Df^*), or *stai* mutant with pan-neuronal expression of *mask* RNAi. Larval NMJs were immunostained with anti-DVGlut (green), anti-DLG (red), and anti-HRP (blue). Arrows point to synaptic boutons that have postsynaptic DLG staining but lack presynaptic DVGLut staining (so-called footprint). Brackets highlight the lateral axons that shows residual DVGlut staining. Scale bars, 10 μm for C; 5 μm for D. (**E**) Quantification of the number of foot-printing boutons in muscle 4 NMJs at segment A3 and A4, as well as axonal accumulation of DVGlut.

### Structure and function analysis of Mask for its action in modulating MT stability

The inclusion of numerous functional domains/motifs in Mask suggests a structural division for its diverse functions. In order to determine the domain(s) requirement for the different functions of Mask, we generated UAS-Mask transgenes that carry a GRGG to GDDG mutation in their KH domain (named UAS-Mask-KH-Mut). Recent studies showed that mutating the GXXG loop to GDDG in the KH minimal motif reduced the ability of the KH domain to bind RNAs (38). The GXXG loop of Mask resides in amino acid 3053-3056 as GRGG, which is completely conserved between fly Mask and human ANKHD1 (corresponding sequence 1710-1713). We also generated UAS-Mask deletion transgenes that lack either N- or C-terminal portions of the protein (depicted in Fig. 5A). One resulting transgene contains the N-terminal two Ankyrin repeats clusters (named Mask-ANK), and the other resulting transgene lacks the N-terminal portion of Mask but contains the NES, NLS, and KH domains (named Mask-KH-Only).

**Figure 5.**
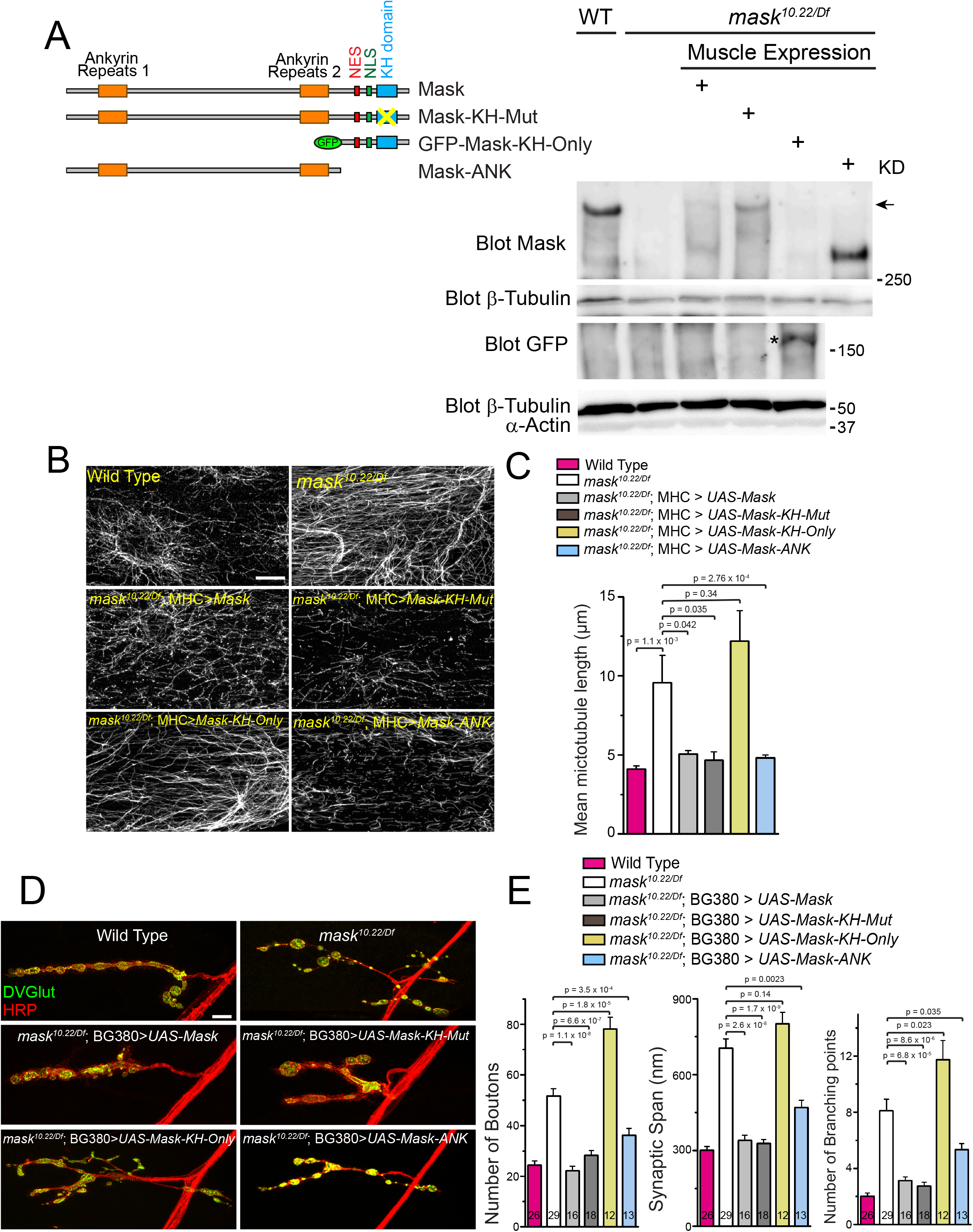
Rescue experiments identify structural requirement for Mask’s action in regulating MT stability. (**A**) A schematic of wild type and mutant UAS-Mask transgenes used in the rescue experiments and the representative western blots showing muscle expression of endogenous Mask in wild type or each of the four UAS-Mask transgenes expressed in muscles in *mask* null mutants. Note that the anti-Mask antibody does not recognize GFP-Mask-KH-Only protein, indicating the antigen of this antibody is outside of the Mask-KH-Only region. Anti-GFP western blots were performed to show the expression of GFP-Mask-KH-Only (indicated by an asterisk). (**B**) Representative confocal images of MT in muscle 6 of wild type, *mask* null (*mask^10.22/Df^)*, and rescues of *mask* null with MHC-Gal4-driven UAS-Mask, UAS-Mask-KH-Mut, UAS-Mask-KH-Only, or UAS-Mask-ANK transgenes. MTs are immunostained with an anti-Acetylated Tubulin antibody. Scale bar: 10 μm. (**C**) Quantification of average MT lengths. (**D**) Representative confocal images of muscle 4 NMJs in wild type, *mask* null (*mask^10.22/Df^*), rescues of *mask* null with pan-neuron (BG380) driven wild type or mutant/deletion UAS-Mask transgenes as shown in A. Scale bar: 10 μm. (**E**) quantification of the number of boutons, synaptic span and the number of branching points at the muscle 4 NMJs. Each data point was normalized to the size of the imaged muscle 4.

We next expressed the mutant *mask* transgenes in the larval muscles in the *mask* null mutants and confirmed that all transgenes express with the predicted molecular weight (Fig. 5A). We then tested the functionality of these transgenes by examining their abilities to rescue the MT-related *mask* mutant defects in larval muscles and NMJs rescue the loss of function phenotypes of Mask in larval muscles and neurons. We found that the UAS-Mask-KH-Mut transgene rescues the MT phenotype in muscles (Fig. 5BC) as well as the defects of NMJ expansion at the NMJs in *mask* mutants (Fig. 5DE) to a level that is comparable to wild type UAS-Mask transgene. These results suggest that the function of the KH domain is not required for the activity of Mask in regulating MT stability or presynaptic terminal expansion. Meanwhile, the UAS-Mask-ANK, but not the UAS-Mask-KH-Only transgene, also rescues the muscle MT length and NMJ overexpansion phenotypes in *mask* mutant larvae (Fig. 5B-E). Together, these data suggest that the N-terminal portion of the Mask protein that contains the Ankyrin repeats domains is sufficient to mediate Mask’s action in regulating MT stability, while the NES, NLS and KH domain are dispensable for the MT-regulating function of Mask.

### Mask inhibits the abundance of the MA-associated protein Jupiter in the axons

To further understand *mask*-mediated regulation of MT stability, we analyzed the distribution and abundance of two MT-associated proteins Jupiter and Futsch proteins in the motor neurons, two other proteins that have been implicated in the stabilization of MT (39, 40). When expressed in the neurons, the Jupiter-mCherry transgene is detected primarily on the axons with low degree extension into the nerve terminals in the area that immediately connects axons and synapses, and these patterns change in response to Mask. The intensity of Jupiter-mCherry in the segmental nerves is substantially enhanced by *mask* loss-of-function, while reduced by Mask overexpression (Fig. 6AB). In addition, the increased abundance of mCherry-Jupiter caused by *mask* RNAi also leads to the extension of mCherry-Jupiter into the nerve terminal; while the reduced mCherry-Jupiter abundance caused by Mask overexpression leads to retraction of mCherry-Jupiter further away from the nerve terminals. To test whether changes of Jupiter-mCherry total protein levels underlie its altered abundances on the MTs, we measured the protein levels of mCherry-Jupiter in larval brain lysates. Neither *mask* RNAi nor overexpression changed the proteins levels of mCherry-Jupiter (Fig. S2A), suggesting that Mask specifically regulates Jupiter’s MT localization which reflects status of MT stability and dynamic. In contrast to Jupiter, we detected no significant changes in Futsch intensity or distribution in segmental nerves or NMJs in response to up- or down-regulation of Mask levels (Fig. 6A-D). Together, these data suggest that Mask negatively regulates the MT localization of MT-associated protein Jupiter. It is worth noting that Jupiter-mCherry and Futsch appear to preferentially label distinct pools of MTs within the axons, which is more prominent in the segmental nerves of Mask-knockdown animals (Fig. S2B). Although there is no precedent evidence supporting the existence of distinct Jupiter-positive and Futsch-positive MT species that may be structurally and functionally different from one another, our data supports the notion that Mask inhibits the decoration of MTs by mCherry-Jupiter but not Futsch.

**Figure 6.**
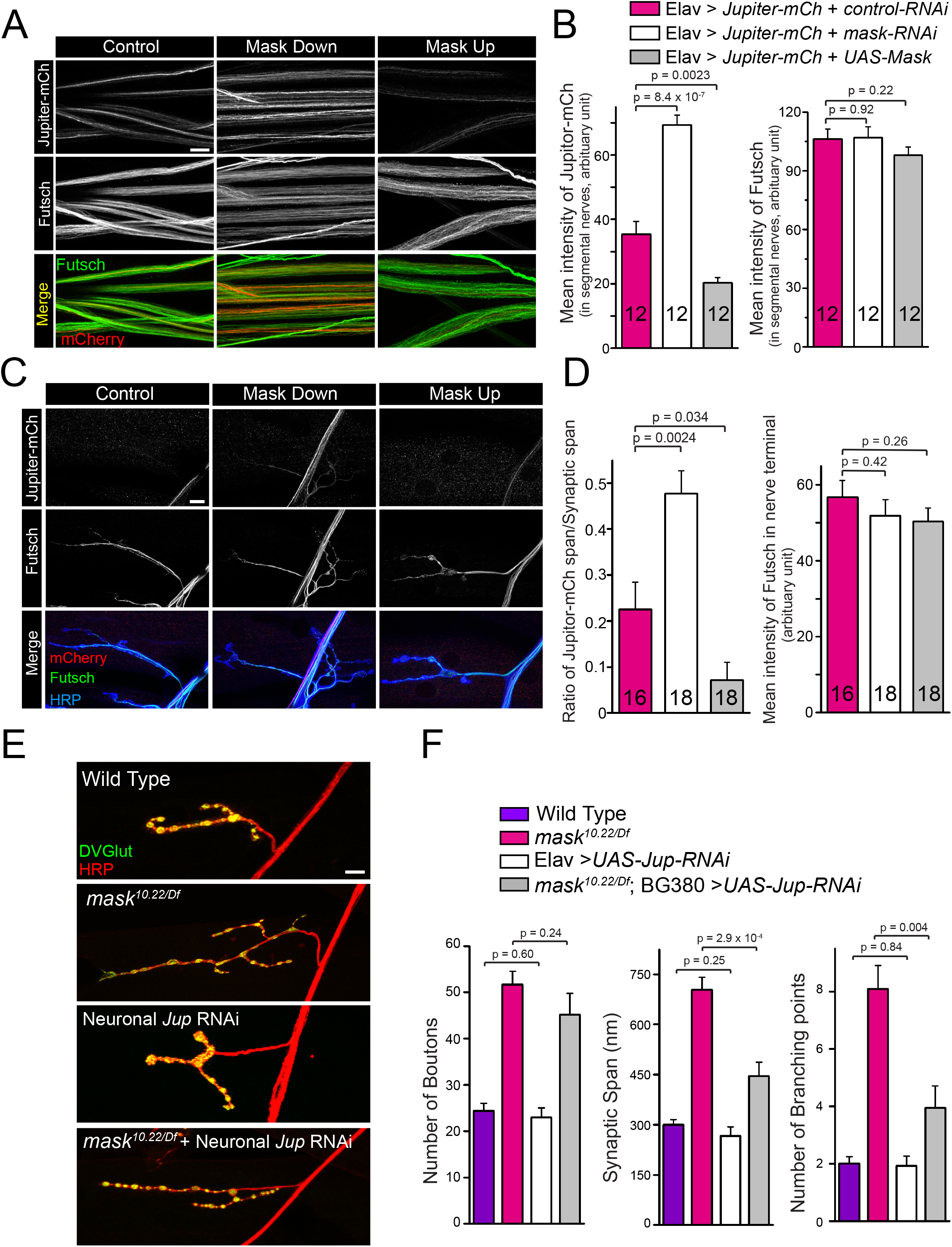
Mask genetically interacts with Jupiter and regulates its localization to microtubules. **(A, C)** Representative confocal images of segmental nerves (A), or muscle 4 NMJs (C), immune-stained with anti-mCherry (Red) and anti-Futsch (green) antibodies in 3^rd^ instar larvae of flies expressing Elav-driven UAS-mCherry-Jupiter together with UAS-control RNAi, *UAS-mask* RNAi or UAS-Mask. **(B, D)** Quantification of the mean intensity of mCherry and Futsch. **(E)** Representative confocal images of muscle 4 NMJs in wild type, *mask* null (*mask^10.22/Df^*), Elav-driven *Jupiter* RNAi (VDRC: KK116151), and BG380-driven *Jupiter* RNAi in the *mask* null mutants. **(F)** Quantification of the number of boutons, synaptic span and the number of branching points at the muscle 4 NMJs. Each data point was normalized to the size of the muscle 4 that were imaged. Scale bars: 10 μm.

Given the molecular regulation of Jupiter by Mask, we next test the functional connections between *jupiter* and *mask* by analyzing the genetic interactions between the two genes. First, we found that overexpressing Jupiter-mCherry in neurons enhances the lethality caused by null mutations of *mask-mask* mutant larvae die at late larval stage; while overexpressing Jupiter in the *mask* null mutant induces lethality at embryonic stage. 26.1% of the progenies (n = 226) from the cross between *mask^10.22^/TM6B* and *mask^Df317^/TM6B* are non-Tubby larvae (*mask^10.22/Df317^)*, and these progenies can survive up to the 3^rd^ instar larval stage. However, no non-Tubby larvae can be found in the progenies (n = 523) when Jupiter is overexpressed in the *mask* mutants. Second, knocking down *Jupiter* in neurons can partially suppress the morphological defects caused by *mask* loss of function at the larval NMJs. A *Jupiter* RNAi line that can efficiently knock down *Jupiter* transcript levels to ~33% of wild type level in neurons (Fig. S3AB), when expressed in *mask* null mutant, can significantly suppress the increased synaptic span and branching point phenotypes (Fig. 6EF) with increased bouton number phenotype in *mask* mutant NMJs unaffected. Together, these strong genetic interactions between *jupiter* and *mask* not only provide a functional link between the two genes but also further support a model where Jupiter works downstream of Mask in regulating MT stability, and that Mask regulates MT dynamics via directly or indirectly inhibiting the MT localization of Jupiter. Furthermore, although Jupiter RNAi does not cause defects in synaptic growth by itself, its genetic interactions with Mask provide new evidence to support the function of Jupiter to stabilize MT.

## Discussion

The characteristics of MT organization and dynamics are cell-type specific. In non-neuronal cells, MTs are labile and undergo frequent growth and shrinkage due to dynamic instability (41). In neurons, MTs are generally more stable than MTs in the dividing cells, however recent studies indicate that there are also abundant labile MT fraction in neurons, especially in the developing neurons, and that a balance between labile and stable MT fractions contribute to normal neuronal functions (4). A key MT-mediated neuronal function is axonal transport. Recent *in vivo* imaging studies of *C. elegans* motor neurons showed that the tiling of the MT fragments in axons is tightly regulated, and such intrinsic MT organization provides the structural basis for efficient cargo transportation along the axons (6).

Decades of studies have demonstrated that MT dynamics are tightly regulated by a number of mechanisms including GTP/GDP ratio, post-translational modification of MT, as well as a vast array of MT-stabilizing, MT polymerizing and depolymerizing, and MT-severing proteins. Our studies identify Mask as a novel regulator of MT stability that controls normal neuronal morphology during development and modulates MT dynamics in concert with factors such as *stai* and Tau that are related to human neuronal pathological conditions. Loss of *mask* resulted in elongated MTs in larval muscles as well as presynaptic over-expansion at the fly larval NMJs, a possible consequence of the over-stabilization of MT given that this phenotype is strongly suppressed by *stai*-induced MT destabilization and partially suppressed by Jupiter knockdown in neurons.

The involvement of Mask in regulating MT stability is further supported by its genetic interactions with *stathmin* and Tau. Previous studies on the effects of overexpressing human Tau protein in fly muscles showed that these ectopic Tau proteins are hyperphosphorylated and cause reduced MT density and enhanced fragmentation, similar to the finding in AD patients and mouse models (16). We found that Mask is able to modulate the Tau-induced MT fragmentation. Loss of Mask enhances, while gain of Mask suppresses MT fragmentation caused by Tau. Neuronal Stathmin family proteins are regulators of MT stability, and perturbation of Stathmin expression impacts neuronal development, plasticity, and regeneration (42). *Drosophila stathmin* mutations cause severely disrupted axonal transport and presynaptic nerve terminal stabilization at the larval NMJs, likely due to impaired integrity of the MT network (12, 13). Knocking out mammalian STMN2, also known as SCG10, results in defects in axon outgrowth and regeneration (15). The result that loss of *mask* in neurons suppresses *stai*-induced axon transport and NMJ development phenotype in a dose-dependent manner, suggesting that Mask antagonizes the action of Stathmin in regulating MT stability. These data also suggested that there is a potential that disturbed dynamics of the MT network under certain pathological conditions can be restored by manipulating Mask levels.

We previously reported that Mask regulates autophagy in fly larval muscles (34). In line with the function of Mask, the levels of ectopic human Tau protein were significantly reduced when co-expressed with Mask, and it is substantially increased when co-expressed with *mask* RNAi (Fig. S1). In addition, Tau proteins in *mask*-knockdown muscles start to form aggregated puncta, possibly due to the excessively elevated levels of the Tau protein. Overexpressing Tau in the muscles causes the developing animals to die at the pupal stages but simultaneously knocking down *mask* suppresses this lethality (unpublished). These findings are very interesting as they demonstrates that the formation of the Tau aggregates does not directly correlate with the toxicity and lethality caused by Tau overexpression in the muscles. It is mostly likely that the toxicity is primarily caused by the severe MT fragmentation induced by Tau expression. MT fragmentation induced by Tau does not seem to be correlative to Tau protein levels, as co-overexpressing Mask with Tau not only significantly reduces the levels of the exogenously expressed Tau protein (Fig. S1) but also potently enhances MT fragmentation (Fig. 1). Although these findings are not directly linked to this study on Mask, they suggest that dysregulation on Tau may cause defects in MT organization that is independent to the toxicity induced by Tau aggregates.

Our loss-of-function studies demonstrated that *mask* is required for normal MT organization in both neurons and muscles. However, overexpression of Mask reduces MT length only under a sensitized background where human Tau induces severe MT fragmentation, but not under an otherwise wild type background, indicating that Mask overexpression itself does not drive significant change in MT dynamics. Mask is a large scaffolding protein containing two ankyrin repeats at its N-terminal and a KH-domain at its C-terminal. Ankyrin-repeats-containing proteins have been implicated in the regulation of MT dynamics. Two isoforms of *Drosophila* Ankyrin2, Ank2-L and Ank2-XL, regulate MT spacing in conjunction with Futsch, and are involved with control of both axon caliber and transport (43). The KH-domain, on the other hand, may mediate Mask’s action in regulating RNA alternative splicing (44) as well as transcription through the HIPPO pathway (20, 21). Our structure/function studies of Mask’s functional domains suggested that the two Ankyrin repeats domains, but not the KH domain, of Mask are essential for Mask’s ability to negatively regulate MT dynamics. The data demonstrate that the Mask transgene containing the N-terminal two Ankyrin repeats domains rescues the MT length in *mask* mutant larval muscles to a level similar to the wild type Mask transgene. Moreover, when expressed in neurons, the same MASK-ANK transgene can also potently rescues synaptic terminal overexpansion in *mask* mutant NMJs, providing not only knowledge for the structural basis of Mask-mediated regulation of MTs but also additional evidence of MT-dependent synaptic regulation by Mask. The sufficiency of Mask’s Ankyrin-repeats domains and the disposableness of the KH domain for MT-regulating function of Mask suggests that protein-protein interactions, but not directly binding to RNAs or DNAs are essential for Mask to regulate MT dynamics. It is interesting that while the KH domain may not be essential for Mask’s action in regulating MT stability, it is required for Mask’s activity in promoting autophagy. The KH domain mutation that diminishes its RNA-binding capacity largely abolishes Mask’s ability to elevate autophagic flux in larval muscles (unpublished data). These results indicate that different functions of Mask are structurally separable and could be modulated independently. Future studies that further investigate the related molecular details will help us to understand how diverse functions of Mask are regulated.

Studies at the fly larval neuromuscular junction have demonstrated complexed mechanisms regulating presynaptic development. A number of evolutionarily conserved pathways, such as the RPM-1/Hiw/Phr1-mediated ubiquitin pathways (45), the BMP signaling pathway (46, 47) and the Wnt signaling pathway (48) are shown to control the normal presynaptic terminal size. Microtubule cytoskeleton also plays essential roles to sustain synaptic morphology and function. Factors that directly regulate microtubule stability have been shown to impact axon transportation or synaptic structures. For example, Stathmin (12, 13), Ringer and Futsch (40), and dTACC (transforming acidic coiled coil) (36) regulate synaptic terminal growth at the fly NMJs, providing direct evidence supporting a link between MT network dynamics and the presynaptic size and morphology. Some signaling pathways control synaptic growth by targeting MT stability, such as the *wg* signaling (49) and the FoxO pathway (50). Our data demonstrate that loss of *mask* function stabilizes MT and promotes presynaptic expansion, providing additional evidence for a strong association between MT stability and synaptic size. Although we don’t have evidence to link Mask to the known signaling pathways that control synaptic growth of the larval NMJs, our imaging and genetic analyses of *mask* loss-of-function phenotypes suggest that Mask negatively regulates MT stability by inhibiting the abundance of Jupiter-associated, but not Futsch-associated MTs. Jupiter is an known microtubule-associated proteins, and has been widely used as a marker for stabilized MTs(39, 51). It’s been speculated that Jupiter promotes MT stabilization, although direct evidence is lacking. Our analysis on Mask and Jupiter here provides additional evidence to support a MT stabilizer function for Jupiter. It is unclear how Mask inhibits MT localization of Jupiter, directly or indirectly, at what cellular compartment, and during what stage of cellular differentiation does this inhibition take place. Mask is a protein that is distributed uniformly in the cytoplasm{Smith, 2002 #737}{Zhu, 2015 #736}, while Jupiter is a MT-associated protein that also shows dynamic localization in embryonic dividing cells{Karpova, 2006 #993}. Could these two proteins interact transiently which impacts the ability of Jupiter to localize to the MTs, or alternatively, Mask may affect properties of MTs which then consequentially prevent Jupiter from binding to MTs? Moreover, Mask may also act at different stages to regulate MT morphology. The result that *mask* genetic null mutants and *mask* RNAi cause MT defects in larval muscles with different severities-the genetic mutants that abolishes Mask functions early and causes more severe phenotypes, while muscle RNAi only diminishes Mask levels at later stage causes milder defects, suggests that Mask regulates MT stability and dynamics at different stages of MT network formation. Although our result suggests that Mask regulates MT through inhibiting MT localization of Jupiter, the result that knocking down Jupiter can only partially suppress *mask* loss of function phenotypes suggests that Jupiter may not be the only downstream effectors of Mask, and other factors may exist to work together with Jupiter to mediate Mask’s effects on MT dynamics. Do these factors actually exist and if yes, what are their molecular identities? All these questions await further investigations.

## Materials and Methods

### Drosophila strains, transgenes and genetics

Flies were maintained at 25°C on standard food. The following strains were used in this study: *mask^10.22^* (25), *mask^Df317^*, UAS-Jupiter-mCherry (52), BG380-Gal4 (neuron-specific) (53), Elav-Gal4 (neuron-specific), MHC-Gal4 (muscle-specific), 24B-GAL4 (muscle-specific), DA-Gal4 (Ubiquitous), UAS-control-RNAi (P{TRiP.JF01147}), UAS-*mask*-RNAi (P{TRiP.HMS01045}) from the Bloomington stock center; UAS-*Jupiter*-RNAi (KK116151) from VDRC. Generation of the full-length wild type *mask* cDNA was previously described (33). This cDNA was used to generate mutant and truncated *mask* cDNA, including pUAST-Mask-KH, pUAST-GFP-Mask-KH-Only, pUAST-Mask-Ank. To generate the UAS-Mask-KH-Mut transgenes, a complementary primer pair containing the mutated coding sequence (GGA<u>GACGAT</u>GGA, mutations underlined) were used to facilitate the change the amino acid sequence (AA3053-3056) from GRGG in the wild type Mask to GDDG in the Mask-KH-Mut transgene. We used the QuickChange II XL site-mutagenesis kit (from Agilent Technology, Santa Clara, CA) to substitute wild type sequence with the mutant sequence. The entire UAS-Mask-KH-Mut coding region was then sequenced to ensure no unintended mutations were introduced. To generate the UAS-Mask-ANK and UAS-Mask-KH-Only transgenes, the KpnI (8001) site was used to separate the Mask encoding region into an N- and a C-terminal fragment, each was then used to generate the pUAST-Mask-ANK and pUAST-Mask-KH-Only respectively. All transgenic fly lines were generated by BestGene Inc. (Chino Hills, CA, USA).

### Western blots

Western blots were performed according to standard procedures. The following primary antibodies were used: mouse anti-β-Tubulin (1:1,000, E7) mouse anti-Alpha-Actin (1:1000, JLA20) from Developmental Studies Hybridoma Bank; rabbit anti-Mask (1:2,000) (25), mouse anti-mCherry antibody (1:1000, 1C51, NBP1-96752 Novus Biologicals), and rabbit anti-GFP (1:1000, A11122, Invitrogen). All secondary antibodies were used at 1:10,000. Data were collected using Luminescent Image Analyzer LAS-3000 (FUJIFILM) and quantified using Multi Gauge (FUJIFILM).

### Immunocytochemistry

Third instar larvae were dissected in ice-cold PBS and fixed in 4% PFA for 30 min. The fixed tissues were stained following standard procedures. The primary antibodies used were: mouse anti-DLG, anti-Futsch and anti-β-Tubulin antibodies from Developmental Studies Hybridoma Bank; rabbit anti-DVGlut (54), rabbit anti-GFP (A11122, Invitrogen) at 1:1000, rabbit anti-mCherry (632496, Clontech) at 1:1000, mouse anti-Acetylated Tubulin (T6793, Sigma) 1:1,000, mouse anti-Tau (12-6400, Invitrogen) at 1: 1000, Cy3-conjugated goat anti-HRP, and Cy5-conjugated goat anti-HRP. The following secondary antibodies (from Jackson ImmunoResearch) were used: Cy3-conjugated goat anti-rabbit IgG at 1:1000, Dylight-488-conjugated anti-mouse IgG at 1:1000, and Alexa-Fluor-647-conjugated goat anti-HRP at 1:1000.

### Confocal imaging and analysis

Single-layer or z-stack confocal images were captured on a Nikon (Tokyo, Japan) C1 confocal microscope. Images shown in the same figure were acquired using the same gain from samples that had been simultaneously fixed and stained. For quantification of microtubule length, z-stack confocal images of microtubule in larval muscle 6/7 in segment A2 were obtained double-blinded, and IMARIS software (Bitplane, Inc) was used to quantify average MT length in randomly-selected muscle areas (see below for details).

The number of boutons and branching points on larval muscle 4 of segments A2-A4 were manually counted. Measurements of synaptic span (total length of NMJs on the muscle surface) as well as the bouton and branching point counts on the same muscle 4 were performed using NIS-Elements (Nikon) imaging software. The synaptic expansions were measured using Nikon elements manual measurement tools. Muscle areas of each corresponding muscle 4 were measured using an eyepiece reticle—a crossline micrometer ruler— under a 20X objective, and the muscle area was used to normalize all three NMJ parameters.

The ration of the nerve terminal distribution of Jupiter-mCherry at the NMJs was measured as the ration of the length of detectable Jupiter-mCherry signal in the nerve terminal to the total length of the same nerve terminal analyzed.

### Quantification of microtubule length in fly larval muscles

Microtubule length was analyzed using IMARIS 9.2.1 imaging software. The analysis was performed in a double-blinded manner. An 80 μm × 80 μm (1024×1024 pixel resolution) confocal image for muscle 6 was used in the analysis, and a vertical step size of 0.15 μm was utilized in order to precisely capture the entire depth of muscle volume containing the microtubule network. From each image, two 30 × 30 μm areas of reasonable clarity were selected for quantification. For each randomly chosen area, 100 +/- 50 tracings were performed manually (assisted by IMARIS), with the total number of traces for each muscle sample averaging ~250 filaments. In order to ensure accurate microtubule tracing and end-to-end length measurements in three-dimensional space, it was necessary to regularly switch between the 3D and layer-by-layer views of the imaged microtubule mesh. Additionally, the angle of the confocal image’s perspective was often changed to allow for further confirmation that a single microtubule filament was being traced.

### Fractionation of Tubulin

Fractionation of β-tubulin was performed as described by Xiong *et al*. with minor modifications (16). Larval muscles of 20 larvae from each genotype were dissected in PBS at room temperature (RT). These muscles were immediately homogenized in 300 ul lysis buffer (150mM KCl, 2mM MgCl2, 50mM Tris, pH 7.5, 2mM EGTA, 2% glycerol, 0.125% Triton X-100, protease inhibitor cocktail) containing either 100 μM or 100 nM Taxol. After incubating for 10 min at RT, the homogenates were centrifuged at 1,500 X g for 5 minutes at RT to remove cellular debris. A small aliquot of the supernatant of each sample was collected to analyze the total β-Tubulin level. The remaining supernatant was ultra-centrifuged at 100,000 g for 30 minutes. After ultracentrifugation, each supernatant and pellet were separated and analyzed by SDS-PAGE and western blots.

### Statistical analysis

Statistical analysis was performed, and graphs were generated in Origin (Origin Lab, Northampton, MA). Each data set was normalized and then compared with other samples in the group (more than two) using ANOVA followed by post-hoc analysis with Turkey’s test, or with the other sample in a group of two using T-test. All histograms are shown as mean ± SEM. The n numbers of each statistical analysis are indicated in each graph.

## Author Contribution

D. M. and M.Z. collected and analyzed the majority of the data. M.Z., C.W. and X.T. designed the experiments. J.J.G. and N.M. provided assistance with data analysis, while H.M. and S.Y. provided technical supports. X.T. and C.W. prepared the manuscript.

## Acknowledgments

We would like to thank Michael Simon for the *mask* mutant alleles and the anti-Mask antibodies, Chris Doe for the UAS-Jupiter-mCherry transgene, the VDRC and the Bloomington Stock center for other fly stocks, and Developmental Studies Hybridoma Bank, created by the NICHD of the NIH and maintained at The University of Iowa, Department of Biology, Iowa City, IA 52242, for antibodies. We also thank Liz McGehee for editorial assistance. This work is supported by a Start-up fund from LSUHSC to X.T. and an ALS Association Grant (18-IIA-414) to C.W.

## Figure Legends

**Supplemental Figure 1.**
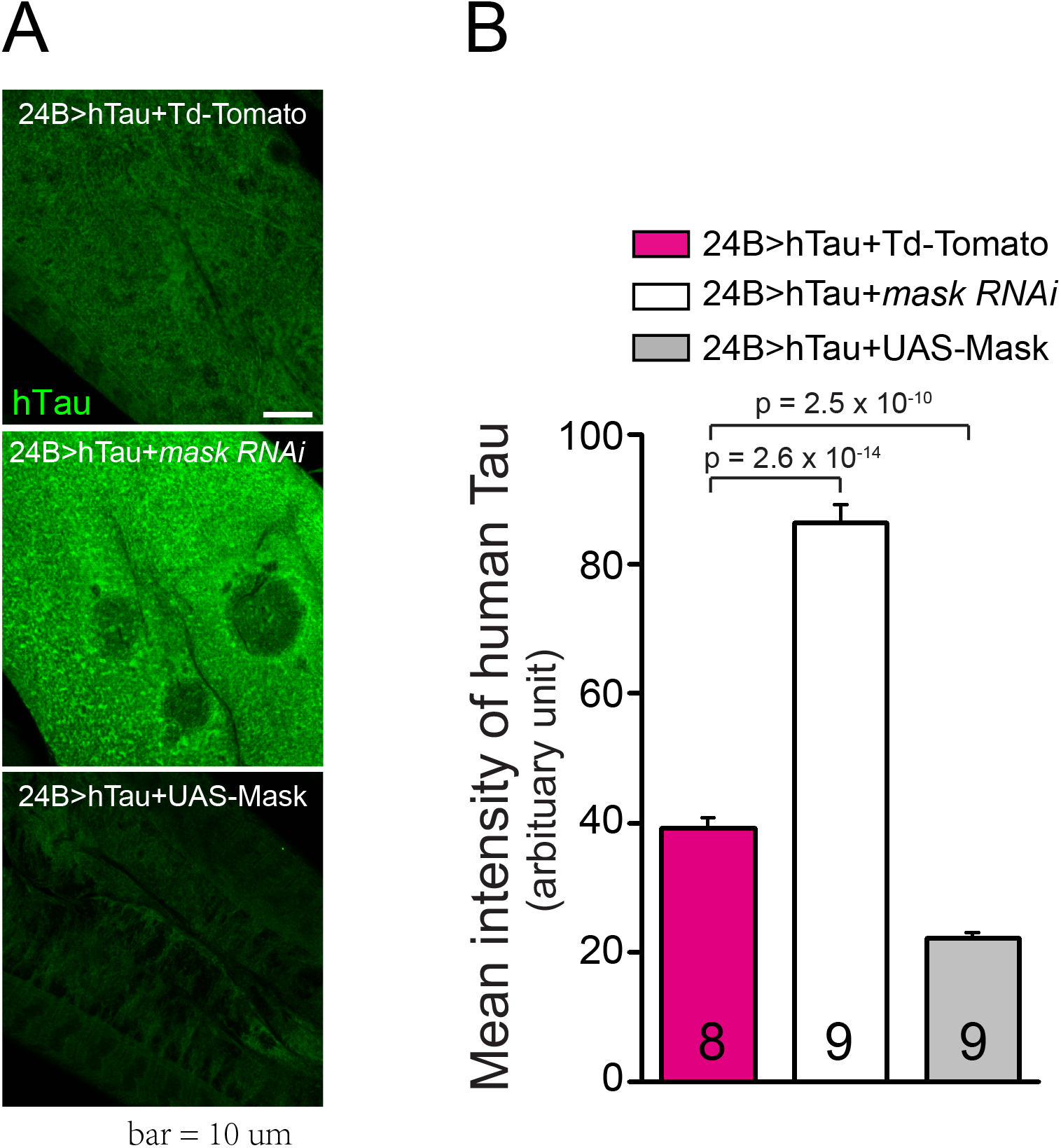
Mask regulates the abundance of Tau protein in larval muscles. (**A**) Representative confocal images of hTau expression in muscle 6 of 24B-Gal4-driven UAS-human Tau (hTau) together with UAS-Td-Tomato, *UAS-mask* RNAi, or UAS-Mask. Human Tau proteins were immunostained with an anti-Tau antibody. Scale bar: 10 μm. (**B**) Quantification of the mean intensity of hTau in the larval muscles.

**Supplemental Figure 2.**
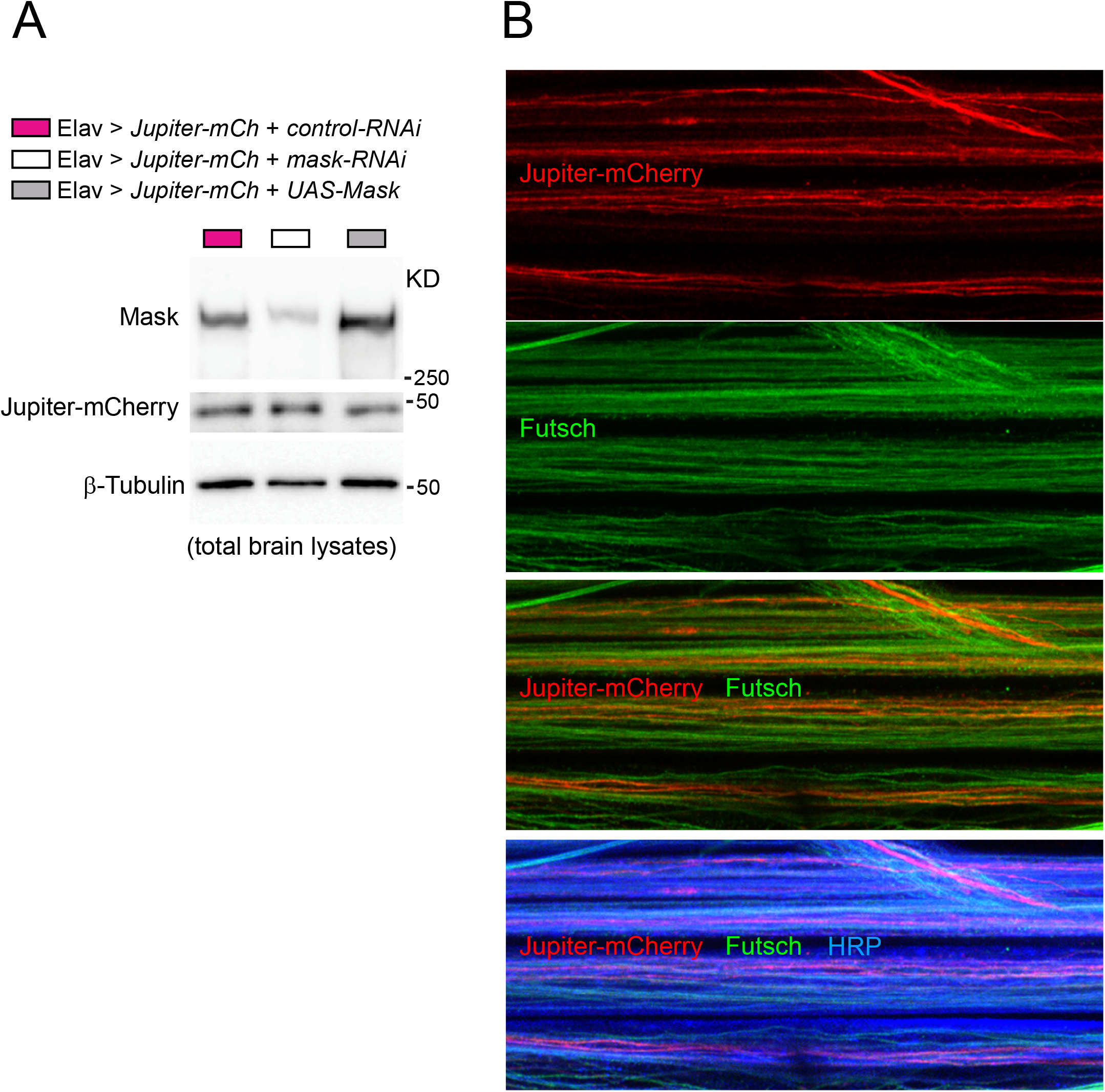
Up- or down-regulation of Mask in fly brains does not change the levels of Jupiter-mCherry expression, and Jupiter-mCherry and Futsch show preferential association to distinct pools of MTs. (**A**) Western blot analysis of larval brains expressing Jupiter-mCherry together with UAS-control-RNAi, UAS-*mask*-RNAi, or UAS-Mask with anti-Mask, anti-mCherry, and anti-β-Tubulin antibodies. (**B**) Representative confocal images of segmental nerves of 3^rd^ instar larvae of Elav-driven Jupiter-mCherry and UAS-*mask*-RNAi. Notice the largely non-overlapping pattern of Jupiter-mCherry and Futsch. Scale bar: 10 μm.

**Supplemental Figure 3.**
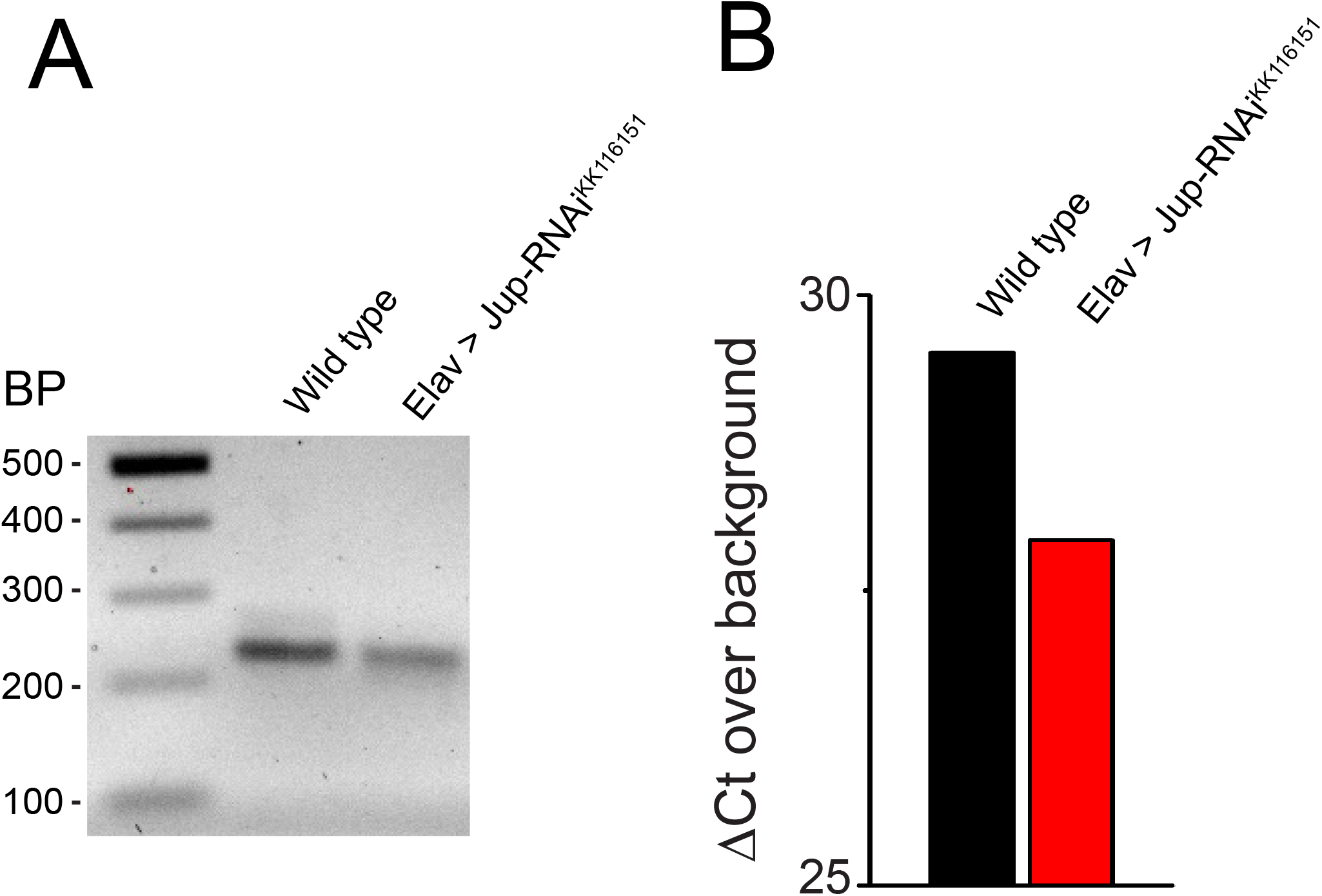
neuronal expression of Jup-RNAi line KK116151 effectively knocks down Jupiter mRNA levels. (**A**) A representative conventional RT-PCR result showing a ~230-bp *Jupiter* PCR fragment amplified from wild type larval brains or brains expressing *UAS-Jup-RNAi^KK116151^* (driven by elav-Gal4). (**B**) Quantitative RT-PCR shows reduction of Jupiter mRNA to ~33% (2^-ΔΔCt) compared to wild type control. The transcript of RPL32was used as the internal control (reference gene) to calculate the ΔCt in control and RNAi samples. Note that the y axis values are expressed as log2 values relative to levels in wells containing no cDNA (“backbround”=Ct 40) following RPL32 normalization; hence, zero indicates essentially no detectable levels of given amplicon and higher numbers = higher amounts of starting RNA.

## Notes

### Competing Interest Statement

The authors have declared no competing interest.

## References

1. Kapitein LC, Hoogenraad CC. Building the Neuronal Microtubule Cytoskeleton. Neuron. 2015;87(3):492–506.

2. van der Vaart B, Akhmanova A, Straube A. Regulation of microtubule dynamic instability. Biochem Soc Trans. 2009;37(Pt 5):1007–13.

3. Hur EM, Saijilafu, Zhou FQ. Growing the growth cone: remodeling the cytoskeleton to promote axon regeneration. Trends Neurosci. 2012;35(3):164–74.

4. Baas PW, Rao AN, Matamoros AJ, Leo L. Stability properties of neuronal microtubules. Cytoskeleton (Hoboken, NJ). 2016;73(9):442–60.

5. Hahn I, Voelzmann A, Liew Y-T, Costa-Gomes B, Prokop A. The model of local axon homeostasis - explaining the role and regulation of microtubule bundles in axon maintenance and pathology. Neural Development. 2019;14(1):11.

6. Yogev S, Cooper R, Fetter R, Horowitz M, Shen K. Microtubule Organization Determines Axonal Transport Dynamics. Neuron. 2016;92(2):449–60.

7. Bowne-Anderson H, Hibbel A, Howard J. Regulation of Microtubule Growth and Catastrophe: Unifying Theory and Experiment. Trends in cell biology. 2015;25(12):769–79.

8. Rossi G, Redaelli V, Contiero P, Fabiano S, Tagliabue G, Perego P, et al. Tau Mutations Serve as a Novel Risk Factor for Cancer. Cancer research. 2018;78(13):3731–9.

9. Tararuk T, Ostman N, Li W, Bjorkblom B, Padzik A, Zdrojewska J, et al. JNK1 phosphorylation of SCG10 determines microtubule dynamics and axodendritic length. J Cell Biol. 2006;173(2):265–77.

10. Wen HL, Lin YT, Ting CH, Lin-Chao S, Li H, Hsieh-Li HM. Stathmin, a microtubule-destabilizing protein, is dysregulated in spinal muscular atrophy. Human molecular genetics. 2010;19(9):1766–78.

11. Strang KH, Golde TE, Giasson BI. MAPT mutations, tauopathy, and mechanisms of neurodegeneration. Laboratory investigation; a journal of technical methods and pathology. 2019;99(7):912–28.

12. Graf ER, Heerssen HM, Wright CM, Davis GW, DiAntonio A. Stathmin is required for stability of the Drosophila neuromuscular junction. J Neurosci. 2011;31(42):15026–34.

13. Duncan JE, Lytle NK, Zuniga A, Goldstein LS. The Microtubule Regulatory Protein Stathmin Is Required to Maintain the Integrity of Axonal Microtubules in Drosophila. PloS one. 2013;8(6):e68324.

14. Liedtke W, Leman EE, Fyffe RE, Raine CS, Schubart UK. Stathmin-deficient mice develop an age-dependent axonopathy of the central and peripheral nervous systems. The American journal of pathology. 2002;160(2):469–80.

15. Klim JR, Williams LA, Limone F, Guerra San Juan I, Davis-Dusenbery BN, Mordes DA, et al. ALS-implicated protein TDP-43 sustains levels of STMN2, a mediator of motor neuron growth and repair. Nat Neurosci. 2019;22(2):167–79.

16. Xiong Y, Zhao K, Wu J, Xu Z, Jin S, Zhang YQ. HDAC6 mutations rescue human tau-induced microtubule defects in Drosophila. Proc Natl Acad Sci U S A. 2013;110(12):4604–9.

17. Braak H, Braak E. Neuropathological stageing of Alzheimer-related changes. Acta neuropathologica. 1991;82(4):239–59.

18. Wittmann CW, Wszolek MF, Shulman JM, Salvaterra PM, Lewis J, Hutton M, et al. Tauopathy in Drosophila: neurodegeneration without neurofibrillary tangles. Science. 2001;293(5530):711–4.

19. Mosavi LK, Cammett TJ, Desrosiers DC, Peng ZY. The ankyrin repeat as molecular architecture for protein recognition. Protein science: a publication of the Protein Society. 2004;13(6):1435–48.

20. Sidor CM, Brain R, Thompson BJ. Mask proteins are cofactors of Yorkie/YAP in the Hippo pathway. Current biology: CB. 2013;23(3):223–8.

21. Sansores-Garcia L, Atkins M, Moya IM, Shahmoradgoli M, Tao C, Mills GB, et al. Mask is required for the activity of the Hippo pathway effector Yki/YAP. Current biology: CB. 2013;23(3):229–35.

22. Musco G, Kharrat A, Stier G, Fraternali F, Gibson TJ, Nilges M, et al. The solution structure of the first KH domain of FMR1, the protein responsible for the fragile X syndrome. Nature structural biology. 1997;4(9):712–6.

23. Garcia-Mayoral MF, Hollingworth D, Masino L, Diaz-Moreno I, Kelly G, Gherzi R, et al. The structure of the C-terminal KH domains of KSRP reveals a noncanonical motif important for mRNA degradation. Structure (London, England: 1993). 2007;15(4):485–98.

24. Grishin NV. KH domain: one motif, two folds. Nucleic acids research. 2001;29(3):638–43.

25. Smith RK, Carroll PM, Allard JD, Simon MA. MASK, a large ankyrin repeat and KH domain-containing protein involved in Drosophila receptor tyrosine kinase signaling. Development (Cambridge, England). 2002;129(1):71–82.

26. Kallappagoudar S, Varma P, Pathak RU, Senthilkumar R, Mishra RK. Nuclear matrix proteome analysis of Drosophila melanogaster. Mol Cell Proteomics. 2010;9(9):2005–18.

27. Müller H, Schmidt D, Steinbrink S, Mirgorodskaya E, Lehmann V, Habermann K, et al. Proteomic and functional analysis of the mitotic Drosophila centrosome. The EMBO journal. 2010;29(19):3344–57.

28. Traina F, Favaro PM, Medina Sde S, Duarte Ada S, Winnischofer SM, Costa FF, et al. ANKHD1, ankyrin repeat and KH domain containing 1, is overexpressed in acute leukemias and is associated with SHP2 in K562 cells. Biochimica et biophysica acta. 2006;1762(9):828–34.

29. Dhyani A, Duarte AS, Machado-Neto JA, Favaro P, Ortega MM, Olalla Saad ST. ANKHD1 regulates cell cycle progression and proliferation in multiple myeloma cells. FEBS letters. 2012;586(24):4311–8.

30. Machado-Neto JA, Lazarini M, Favaro P, Franchi GC, Jr., Nowill AE, Saad ST, et al. ANKHD1, a novel component of the Hippo signaling pathway, promotes YAP1 activation and cell cycle progression in prostate cancer cells. Experimental cell research. 2014;324(2):137–45.

31. Dhyani A, Machado-Neto JA, Favaro P, Saad ST. ANKHD1 represses p21 (WAF1/CIP1) promoter and promotes multiple myeloma cell growth. European journal of cancer (Oxford, England: 1990). 2015;51(2):252–9.

32. Machado-Neto JA, Lazarini M, Favaro P, de Melo Campos P, Scopim-Ribeiro R, Franchi Junior GC, et al. ANKHD1 silencing inhibits Stathmin 1 activity, cell proliferation and migration of leukemia cells. Biochimica et biophysica acta. 2015;1853(3):583–93.

33. Zhu M, Li X, Tian X, Wu C. Mask loss-of-function rescues mitochondrial impairment and muscle degeneration of Drosophila pink1 and parkin mutants. Human molecular genetics. 2015;24(11):3272–85.

34. Zhu M, Zhang S, Tian X, Wu C. Mask mitigates MAPT- and FUS-induced degeneration by enhancing autophagy through lysosomal acidification. Autophagy. 2017;13(11):1924–38.

35. Zhu M, Zhang S, Tian X, Wu C. Mask mitigates MAPT- and FUS-induced degeneration by enhancing autophagy through lysosomal acidification. Autophagy. 2017:1–15.

36. Chou VT, Johnson S, Long J, Vounatsos M, Van Vactor D. dTACC restricts bouton addition and regulates microtubule organization at the Drosophila neuromuscular junction. Cytoskeleton (Hoboken, NJ). 2020;77(1-2):4–15.

37. Akhmanova A, Steinmetz MO. Control of microtubule organization and dynamics: two ends in the limelight. Nature reviews Molecular cell biology. 2015;16(12):711–26.

38. Hollingworth D, Candel AM, Nicastro G, Martin SR, Briata P, Gherzi R, et al. KH domains with impaired nucleic acid binding as a tool for functional analysis. Nucleic acids research. 2012;40(14):6873–86.

39. Karpova N, Bobinnec Y, Fouix S, Huitorel P, Debec A. Jupiter, a new Drosophila protein associated with microtubules. Cell motility and the cytoskeleton. 2006;63(5):301–12.

40. Shi Q, Lin YQ, Saliba A, Xie J, Neely GG, Banerjee S. Tubulin Polymerization Promoting Protein, Ringmaker, and MAP1B Homolog Futsch Coordinate Microtubule Organization and Synaptic Growth. Frontiers in cellular neuroscience. 2019;13:192–.

41. Mitchison T, Kirschner M. Dynamic instability of microtubule growth. Nature. 1984;312(5991):237–42.

42. Chauvin S, Sobel A. Neuronal stathmins: a family of phosphoproteins cooperating for neuronal development, plasticity and regeneration. Progress in neurobiology. 2015;126:1–18.

43. Stephan R, Goellner B, Moreno E, Frank CA, Hugenschmidt T, Genoud C, et al. Hierarchical Microtubule Organization Controls Axon Caliber and Transport and Determines Synaptic Structure and Stability. Developmental cell. 2015;33(1):5–21.

44. Brooks AN, Duff MO, May G, Yang L, Bolisetty M, Landolin J, et al. Regulation of alternative splicing in Drosophila by 56 RNA binding proteins. Genome research. 2015;25(11):1771–80.

45. Tian X, Wu C. The role of ubiquitin-mediated pathways in regulating synaptic development, axonal degeneration and regeneration: insights from fly and worm. The Journal of physiology. 2013;591(Pt 13):3133–43.

46. Keshishian H, Kim YS. Orchestrating development and function: retrograde BMP signaling in the Drosophila nervous system. Trends Neurosci. 2004;27(3):143–7.

47. Bayat V, Jaiswal M, Bellen HJ. The BMP signaling pathway at the Drosophila neuromuscular junction and its links to neurodegenerative diseases. Curr Opin Neurobiol. 2011;21(1):182–8.

48. Koles K, Budnik V. Wnt signaling in neuromuscular junction development. Cold Spring Harbor perspectives in biology. 2012;4(6).

49. Franco B, Bogdanik L, Bobinnec Y, Debec A, Bockaert J, Parmentier ML, et al. Shaggy, the homolog of glycogen synthase kinase 3, controls neuromuscular junction growth in Drosophila. The Journal of neuroscience: the official journal of the Society for Neuroscience. 2004;24(29):6573–7.

50. Nechipurenko IV, Broihier HT. FoxO limits microtubule stability and is itself negatively regulated by microtubule disruption. The Journal of cell biology. 2012;196(3):345–62.

51. Takeda M, Sami MM, Wang YC. A homeostatic apical microtubule network shortens cells for epithelial folding via a basal polarity shift. Nature cell biology. 2018;20(1):36–45.

52. Chabu C, Doe CQ. Dap160/intersectin binds and activates aPKC to regulate cell polarity and cell cycle progression. Development. 2008;135(16):2739.

53. Budnik V, Koh YH, Guan B, Hartmann B, Hough C, Woods D, et al. Regulation of synapse structure and function by the Drosophila tumor suppressor gene dlg. Neuron. 1996;17(4):627–40.

54. Daniels RW, Collins CA, Gelfand MV, Dant J, Brooks ES, Krantz DE, et al. Increased expression of the Drosophila vesicular glutamate transporter leads to excess glutamate release and a compensatory decrease in quantal content. J Neurosci. 2004;24(46):10466–74.

